# Abnormal pro-inflammatory immune cell responses precede the clinical onset of hypertensive disorders of pregnancy

**DOI:** 10.1101/2025.08.04.668579

**Authors:** Ayman Rezk, Chelsea A DeBolt, Gautier Breville, Rui Li, Valerie A Riis, Liqhwa Ncube, Fatoumata Barry, Michal A Elovitz, Amit Bar-Or

## Abstract

**Background:** Hypertensive disorders of pregnancy (HDP) are significant contributors to maternal morbidity and mortality as well as to long-term cardiovascular health, with an inequitable burden among Black individuals. While there is growing interest in the possibility that inflammatory responses are involved with HDP, work to date has primarily been cross-sectional, and immune cell profiling has focused on cell phenotyping that may not capture relevant functional properties of the immune cells.

**Objective:** To examine whether an inflammatory cellular response is found in the blood of pregnant women preceding the clinical onset of HDP.

**Study design:** This nested case-control study included 27 pregnant women who later developed HDP and 73 who had a healthy pregnancy delivered at term, all of whom provided blood samples during the second trimester.

**Results:** We observed a greater abundance of peripheral blood T cells in individuals later diagnosed with HDP. Among these were CD161^+^ CD4 T cells that showed increased expression of pro-inflammatory cytokines, including GM-CSF and TNF. Similarly, monocytes from these individuals exhibited increased expression of pro-inflammatory cytokines. Anti-inflammatory T cells and monocytes were not altered in those with versus without a future diagnosis of HDP.

**Conclusion:** We identify a pro-inflammatory cellular immune signature in pregnant individuals destined to have HDP. Immune signatures may serve as a new biomarker to identify subsets of individuals at particular risk for HDP and/or point to new therapeutic targets to prevent HDP.

## Introduction

Hypertensive disorders of pregnancy (HDP), including preeclampsia and gestational hypertension, complicate up to 15% of pregnancies with significant perinatal morbidity as well as long-term health risks to the mother ^1–6^. The prevalence of HDP and its subsequent adverse effects on maternal and infant health are higher among historically marginalized groups ^1,5,7–10^. Survivors of HDP face a 2-fold increased risk of future cardiovascular disease compared to normotensive women ^2,11–13^. Women with the most severe form of HDP, preeclampsia with severe features, are at 4 times greater risk of developing heart failure, and 2 times more likely to develop ischemic heart disease and death from cardiovascular causes compared to women without HDP ^2,11–13^. Preeclampsia and eclampsia are also associated with adverse cerebrovascular health later in life, including seizure disorder, stroke, and vascular dementia ^2,11,12,14,15^. Black women are disproportionately affected by diagnoses of HDP and the subsequent adverse effects on maternal and infant health ^1,5,7–10^. Not only do Black women experience a higher rate of HDP, but they are more likely to develop the severest form of HDP, preterm preeclampsia ^10,16^.

Maternal infection was considered a potential etiology of HDP since the start of the 20^th^ century, with the concept that ‘toxins’ mediate the disease ^17^. As HDP lacks traditional features of infection, such as fever, leukocytosis, and/or clinical signs of sepsis, the concept of a pathogenic organism causing HDP is not supported. However, epidemiological research continues to demonstrate associations between maternal infection and HDP; specifically, associations between periodontal disease, urinary tract infections, and COVID-19 with HDP have been reported ^18,19^. A potential immunological mechanism for these associations is the induction of inflammatory pathways from these infections. While immune perturbations occur as part of normal pregnancy ^20,21^, there is evidence of increased inflammation in some cases of HDP ^22^. Further, while a systemic proinflammatory cytokine signature is associated with HDP ^23–25^, changes in immunity also occur locally at the level of the uteroplacental interface, with dysregulation of immune cells such as NK cells, B cells, and macrophages in pregnancies complicated by HDP ^26,27^. Perturbations in T-cell immune responses have also been associated with poor placental invasion, which is considered a critical mechanism underlying the development of HDP ^22^. Alterations in peripheral T cell functions have been associated with HDP, including enhanced pro-inflammatory CD4 T cell responses and diminished regulatory T cell responses at term ^28–31^. Because of the substantial temporal gap between the biological development of HDP and the onset of clinical disease symptoms, immune profiles that are etiologically relevant may no longer be present when HDP is well established. Furthermore, when immune perturbations are detected at the time of established HDP, it is difficult to know whether they contribute to the abnormal process or reflect a consequence, since the immune system would normally be expected to respond to any pathologic process and may appear changed.

To assess whether abnormal immune cell shifts may occur before the clinical recognition of HDP, we studied a prospective cohort of pregnant individuals at higher risk of developing HDP and measured functional immune profiles of circulating cells during the 2nd trimester of pregnancy, then compared the immune profiles of those who subsequently did versus did not develop HDP months later. Our analysis uncovered novel pro-inflammatory cytokine signatures in CD161^+^ T cells and monocytes from individuals who subsequently met clinical diagnostic criteria for HDP.

## Methods

### Study approval

Black pregnant individuals meeting the inclusion criteria were prospectively recruited at the Hospital of the University of Pennsylvania. All participants provided written informed consent. The study was approved by and carried out per the World Medical Association Declaration of Helsinki and the guidelines and regulations of the Institutional Review Board of the University of Pennsylvania (IRB # 842850).

### Study participants

Patient selection is shown in Supplementary Figure 1. 120 individuals were enrolled, with 117 individuals having available blood specimens to analyze. Maternal blood was collected between gestational weeks 16 and 24. Patients with spontaneous preterm birth (n=7), chronic hypertension (n=8), and medically indicated preterm birth for severe fetal growth restriction (n=1) were excluded from the analysis. Of the 100 individuals included, 27 participants were later diagnosed with HDP according to the ACOG definition ^32^, and 73 were not.

### Peripheral blood mononuclear cell isolation and cryopreservation

Whole blood was diluted with PBS containing 2mM EDTA (ThermoFisher) and overlayed on Ficoll (GE Healthcare) in SepMate tubes (StemCell Technologies). Tubes were spun at 1200xg for 10 minutes at room temperature with brakes on. The white cell monolayer was collected and washed twice before cell cryopreservation in human AB serum (Sigma Aldrich) containing 10% DMSO (Fisher Scientific). Cryopreserved cells were stored in the vapor phase of liquid nitrogen.

### Immunophenotyping of whole blood

An aliquot of whole blood was diluted 1:10 in 3% acetic acid with methylene blue (StemCell Technologies) to count white blood cells. Whole blood was lysed in 1X RBC lysis solution (Biologend) for 15 minutes at room temperature. The cell suspension was washed once, and the cell pellet was resuspended in a cocktail of antibodies detailed in Supplementary Tables 1 and 2 (Please see the Major Resources Table in the Supplemental Materials). Cells were incubated at room temperature for 30 minutes and washed once before fixation with BD CytoFix/CytoPerm solution (BD Biosciences). Cells were washed twice before data acquisition on a BD LSRFortessa^TM^ (BD Biosciences). Data analysis was performed using FlowJo (BD Biosciences). Acquisition and analysis were carried out by a blinded operator. The data output was used to calculate the absolute counts of immune cell populations using whole blood cell count.

### Immunophenotyping of cryopreserved PBMC

Cryopreserved vials were thawed in a water bath set at 37℃°C. Cells were diluted in X-VIVO 10 (Lonza) containing DNAe (0.5 μg/ml; StemCell Technologies). Cells were washed once, resuspended in X-VIVO 10 containing DNAse (1μg/ml), and incubated in a humidified incubator for 15 minutes. Cells were washed once, resuspended in X-VIVO 10, and 10^6^ live cells were transferred into a 96-well V-shaped plate overnight in a humidified incubator. Where indicated, cells were stimulated with phorbol 12-myristate 13-acetate (10ng/ml, Sigma Aldrich) and ionomycin (500ng/ml, Sigma Aldrich) in the presence of monensin A (BD Biosciences) for 5 hours in a humidified incubator. Cells were harvested, washed once, and incubated with a PBS solution containing LIVE/DEAD fixable aqua dead cell stain (ThermoFisher Scientific) for 30 minutes at room temperature. Cells were washed once, and the cell pellet was resuspended in the appropriate antibody cocktail detailed in Supplementary Tables 1 and 2 (Please see the Major Resources Table in the Supplemental Materials). Cells were incubated at room temperature for 30 minutes or at 37℃ °C for 15 minutes, followed by room temperature for 15 minutes. Cells were washed once and incubated with BD CytoFix/CytoPerm solution (BD Biosciences) for 30 minutes or FOXP3/Transcriptional Factor staining buffer (ThermoFisher) for 60 minutes at 4℃°C. Cells were washed twice, resuspended in the appropriate wash buffer, and left overnight in a 4°C fridge. Where indicated, cells were incubated with a cocktail of antibodies directed against intracellular protein (Supplementary Tables 1 and 2). Cells were washed twice before data acquisition on a BD LSRFortessa^TM^ (BD Biosciences). Data analysis was performed using FlowJo (BD Biosciences). A blinded operator carried out acquisition and analysis.

### Statistical analysis

Statistical analysis was performed using Prism 10 (GraphPad Software, San Diego, CA). Unpaired data were compared using a non-parametric Mann-Whitney test, followed by correction for multiple comparisons using the Holm-Sidak method. A *p*-value < 0.05 was considered statistically significant.

## Results

### Elevated counts of T cells in the blood of pregnant women with a future diagnosis of HDP

According to the standard American College of Obstetrics and Gynecology (ACOG) definition ^32^, 27 pregnant women were later diagnosed with HDP, while 73 did not receive the diagnosis in our cohort. Basic demographic information of the two groups is summarized in Table 1. While the groups were well balanced for most features, including gestational age at time of blood sampling, individuals who were later diagnosed with HDP had a higher median body mass index (BMI) and were more likely to have pregestational diabetes mellitus (Table 1). Flow cytometric analysis of fresh whole blood collected during the second trimester revealed no significant differences between the two groups in the absolute counts of several immune cell populations including eosinophils, basophils, neutrophils, classical monocytes, intermediate monocytes, non-classical monocytes, CD11c^+^ dendritic cells, CD123^+^ dendritic cells, CD56^bright^ NK cells and CD56^+^CD16^+^ NK cells (Figure 1B). However, total T cell count, including CD4^+^ T cells, was significantly elevated in the group that subsequently received a diagnosis of HDP (Figure 1B).

**Figure 1.**
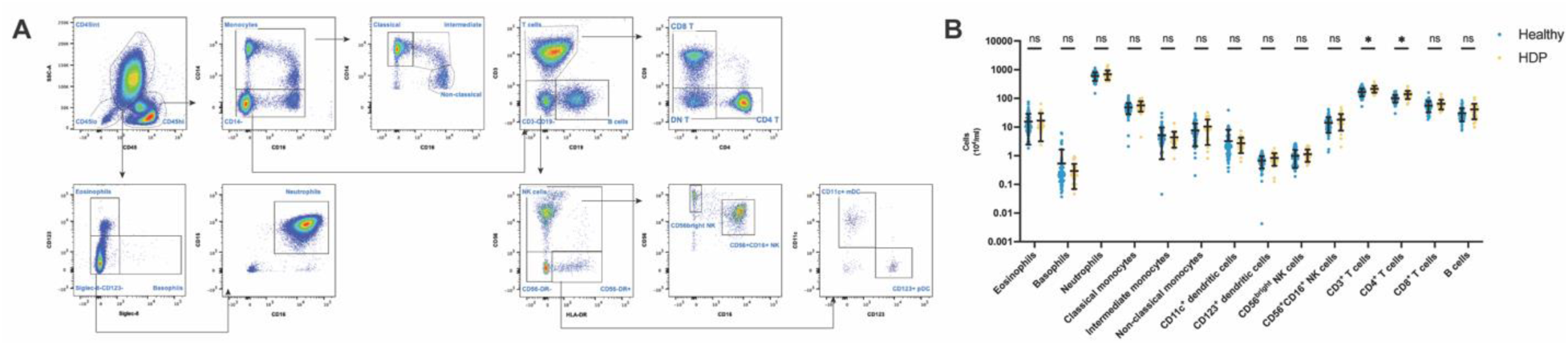
Elevated count of T cells in individuals with a subsequent diagnosis of HDP. (A) Representative gating strategy of immune cells in fresh whole blood. Cells were divided into CD45^int^ and CD45^hi^ gates. CD45^int^ cells were further defined as Siglec-8^+^CD123^-^ (eosinophils), Siglec-8^-^CD123^+^ (basophils), and Siglec-8^-^CD123^-^CD15^+^CD16^+^ (neutrophils). CD45^hi^ cells were divided into CD14^+^CD16^-^ (classical monocytes), CD14^+^CD16^+^ (intermediate monocytes), CD14^lo^CD16^+^ (non-classical monocytes), CD19^+^ (B cells), CD4^+^CD3^+^ (CD4 T cells), CD8^+^CD3^+^ (CD8 T cells), CD56^bright^ and CD56^+^CD16^+^ CD3^-^CD19^-^CD14^-^ (NK cell subsets), CD11c^+^ and CD123^+^ HLA-DR^+^CD56^-^CD3^-^CD19^-^CD14^-^ (dendritic cell subsets). (B) Absolute counts of immune cell populations in the blood of healthy pregnant women and those who later developed HDP. N=73 healthy and n=27 HDP. Data is shown as mean ± SD.

**Table 1.** Basic demographics of the cohort.

|  | **Group 1: Controls** | **Group 2: HDP** | **P-value** |
| --- | --- | --- | --- |
|  | (N=73) | (N=27) |  |
| **Race** |  |  |  |
| Black | 69 (94.5%) | 27 (100%) | 0.505 |
| Mixed/Other/Unknown | 4 (5.5%) | 0 (0%) |  |
| **Ethnicity** |  |  |  |
| Hispanic/Latino | 8 (11.0%) | 1 (3.7%) | 0.464 |
| Non-Hispanic/Non-Latino | 65 (89.0%) | 26 (96.3%) |  |
| **Age at delivery** |  |  |  |
| Median [IQR] | 26.0 [10.0] | 26.0 [8.50] | 0.547 |
| **Education** |  |  |  |
| HS or less | 40 (54.8%) | 16 (59.3%) | 0.891 |
| Post college/degree | 4 (5.5%) | 1 (3.7%) |  |
| Some college/degree | 29 (39.7%) | 10 (37.0%) |  |
| **BMI at 1st prenatal** |  |  |  |
| Median [IQR] | 26.4 [9.86] | 32.6 [12.9] | 0.043 |
| **BMI Groups** |  |  |  |
| <30 | 48 (65.8%) | 12 (44.4%) | 0.089 |
| >=30 | 25 (34.2%) | 15 (55.6%) |  |
| **Gestational age BMI measured** |  |  |  |
| Median [IQR] | 10.0 [1.00] | 10.0 [0] | 0.38 |
| **Insurance** |  |  |  |
| Private/Individual<br>(HMO/PPO) | 15 (20.5%) | 6 (22.2%) | 1 |
| Public Insurance | 58 (79.5%) | 21 (77.8%) |  |
| **Marital status** |  |  |  |
| Married | 9 (12.3%) | 5 (18.5%) | 0.64 |
| Not Married | 64 (87.7%) | 22 (81.5%) |  |
| **Smoking during pregnancy** |  |  |  |
| No | 66 (90.4%) | 24 (88.9%) | 0.044 |
| Yes | 7 (9.6%) | 1 (3.7%) |  |
| Unknown | 0 (0%) | 2 (7.4%) |  |
| **Pregestational Diabetes** |  |  |  |
| No | 73 (100%) | 24 (88.9%) | 0.026 |
| Yes | 0 (0%) | 3 (11.1%) |  |
| **Gestational Diabetes** |  |  |  |
| No | 69 (94.5%) | 25 (92.6%) | 1 |
| Yes | 4 (5.5%) | 2 (7.4%) |  |
| **Gestational age at delivery** |  |  |  |
| Median [IQR] | 39.5 [1.80] | 38.7 [2.18] | 0.055 |
| **Preterm birth** |  |  |  |
| No | 73 (100%) | 25 (92.6%) | 0.122 |
| Yes | 0 (0%) | 2 (7.4%) |  |
| **Birth weight** |  |  |  |
| Mean (SD) | 3250 (454) | 3190 (545) | 0.632 |
| **Infant sex** |  |  |  |
| Female | 37 (50.7%) | 17 (63.0%) | 0.386 |
| Male | 36 (49.3%) | 10 (37.0%) |  |
| **Gestational age at**<br>**Immune Profiling** |  |  |  |
| Median [IQR] | 20.3 [1.14] | 20.1 [0.784] | 0.764 |

### Peripheral pro-inflammatory CD161^+^ CD4 T cell responses are elevated before the clinical diagnosis of HDP

Next, we aimed to characterize the phenotype and functional profile of CD4 and CD8 T cell subsets. Broadly, the proportions of naïve (CCR7^+^CD45RO^-^), central memory (CCR7^+^CD45RO^+^), effector memory (CCR7^-^CD45RO^-^), and terminal effector (CCR7^-^CD45RO^-^) CD4 and CD8 T cells were similar between pregnant women who were and were not subsequently diagnosed with HDP (Supplementary Figure 2). Normal pregnancy was associated with a shift in the balance between pro-inflammatory T cell responses (Th1 and Th17 cells) to Th2 and Treg cells, in favor of the latter populations ^30,31,33^. We found that the absolute count, but not the frequency, of CD161^+^ CD4 T cells was elevated in the blood of pregnant women subsequently diagnosed with HDP (Figure 2A-B). CD161 characterizes a memory CD4 T cell population that produces pro-inflammatory cytokines such as IFN-γ, IL-17, and TNF ^34,35^, and CD161^+^ CD4 T cells have been found to infiltrate the decidua of first-trimester placentas ^36^.

**Figure 2.**
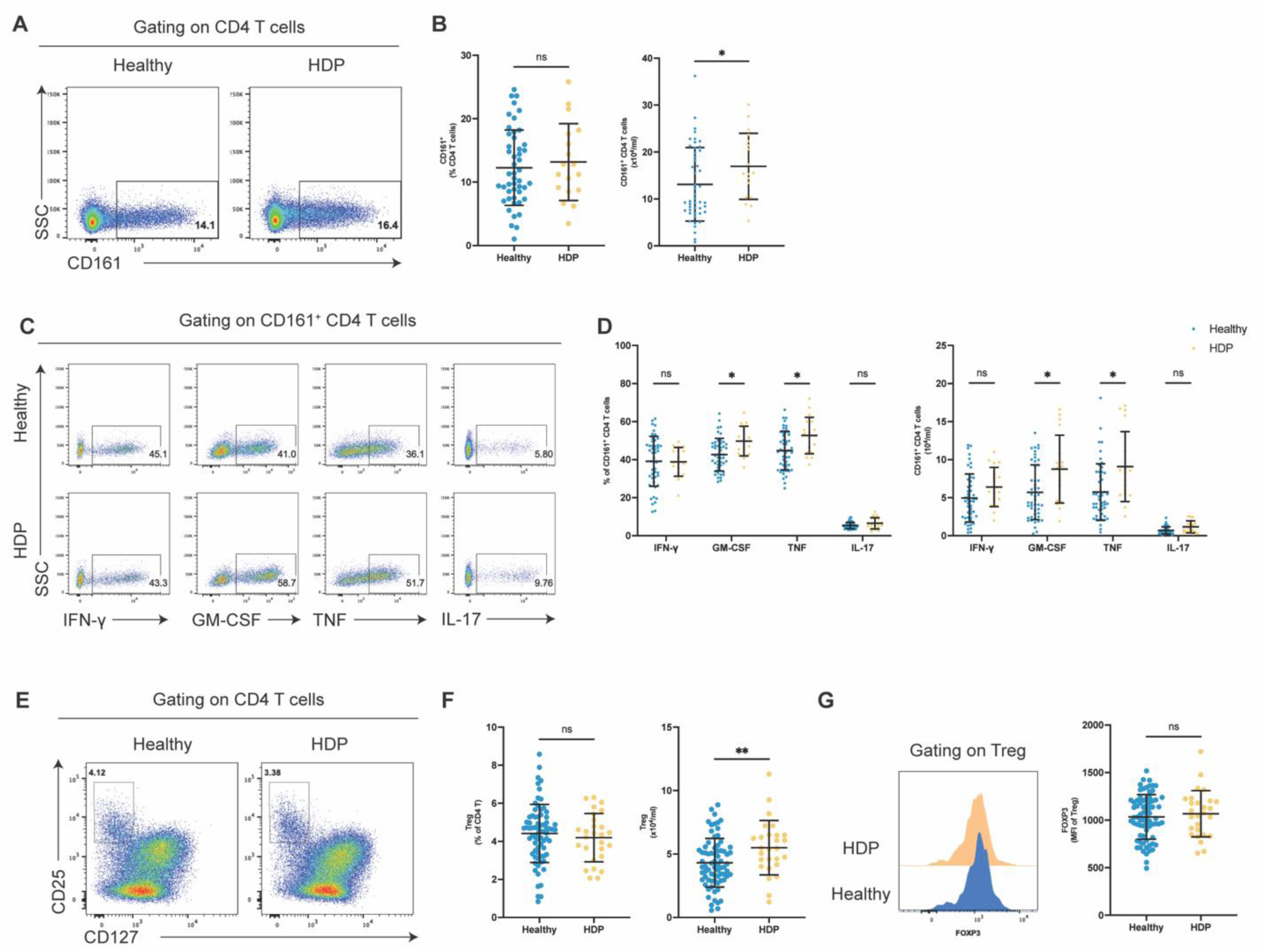
Increased GM-CSF and TNF expression by CD161^+^ CD4 T cells before the clinical diagnosis of HDP. (A) Representative flow cytometry plots showing CD161 expression on CD4 T cells. (B) Frequency and absolute count of CD161^+^ CD4 T cells in the peripheral blood of healthy pregnant women and those who later developed HDP (n=48 healthy and n=18 HDP). (C) Representative flow cytometry plots of IFN-γ and IL-17 expression in CD161^+^ CD4 T cells. (D) Frequencies and absolute counts of cytokine-expressing CD161^+^ CD4 T cells in the peripheral blood of healthy pregnant women and those subsequently diagnosed with HDP (n=48 healthy and n=18 HDP). (E) Representative flow cytometry plots showing CD25^hi^ CD127^lo^ regulatory CD4 T cells among CD4 T cells. (F) Frequency and absolute count of Treg cells in the peripheral blood of healthy pregnant women and those later diagnosed with HDP (n=73 healthy and n=27 HDP). (F) Representative expression of FOXP3 in Treg cells. (G) Summary data showing FOXP3 expression in Treg cells from healthy and HDP pregnancies (n=73 healthy and n=27 HDP). Data is shown as mean ± SD.

Analysis of intracellular cytokine expression following brief activation showed that total CD4 and CD8 T cells did not exhibit noticeable differences in their cytokine expression profile, including IFN-γ, GM-CSF, TNF, IL-17, IL-4, and IL-10, between the two groups (Supplementary Figure 3). However, the frequency and absolute counts of GM-CSF^+^ and TNF^+^ CD161^+^ CD4 T cell subsets were elevated in the HDP group (Figure 2C-D). Finally, the frequency of regulatory CD4 T cells (Tregs), defined as CD25^hi^ CD127^lo^, was similar between the two groups (Figure 2E-F), as was the expression of the master transcriptional factor FOXP3 in the Treg cells (Figure 2G). Altogether, we observed abnormal pro-inflammatory cytokine responses by CD161^+^ CD4 T cells in the peripheral blood of pregnant women that preceded clinical diagnosis of HDP.

### Increased counts of peripheral memory and pro-inflammatory cytokine-expressing B cells before the clinical onset of HDP

Composition of B cells changes throughout normal pregnancy, including reduced frequencies of transitional and activated cells ^33,37^, though this has not been described in the context of HDP risk. We defined B cell sub-populations among total CD19^+^ cells as transitional (CD24^+^CD38^hi^CD10^+^) B cells or as plasmablasts (CD24^-^CD38^+^), as well as either naïve (CD27^-^ IgD^+^), non-class switched memory (CD27^+^IgD^+^), class-switched memory (CD27^+^IgD^-^) or double negative memory (CD27^-^IgD^-^) B cells among the mature (CD24^+^CD38^-^) population (Figure 3A, C). Whilst the frequencies of those populations were similar between the two groups (Figure 3B-D), the absolute counts of non-class and class-switched memory B cells were elevated in the blood of pregnant women with a future diagnosis of HDP (Figure 3E). Expression of CD80 and CD86 (co-stimulatory molecules involved in T cell activation) on unstimulated total B cells was similar between the two groups (Supplementary Figure 4). Among their antibody-independent functions, B cells can express cytokines such as GM-CSF, TNF, IL-6, and IL-10, which are involved in B cell modulation of T cell and myeloid cell responses ^38^. We observed increased counts of B cells expressing GM-CSF^+^ and TNF^+^ (but not IL-10^+^) in pregnant women later diagnosed with HDP (Supplementary Figure 4). Together, these findings indicate that levels of memory and pro-inflammatory B cells are elevated in peripheral blood before diagnosing HDP in pregnant women.

**Figure 3.**
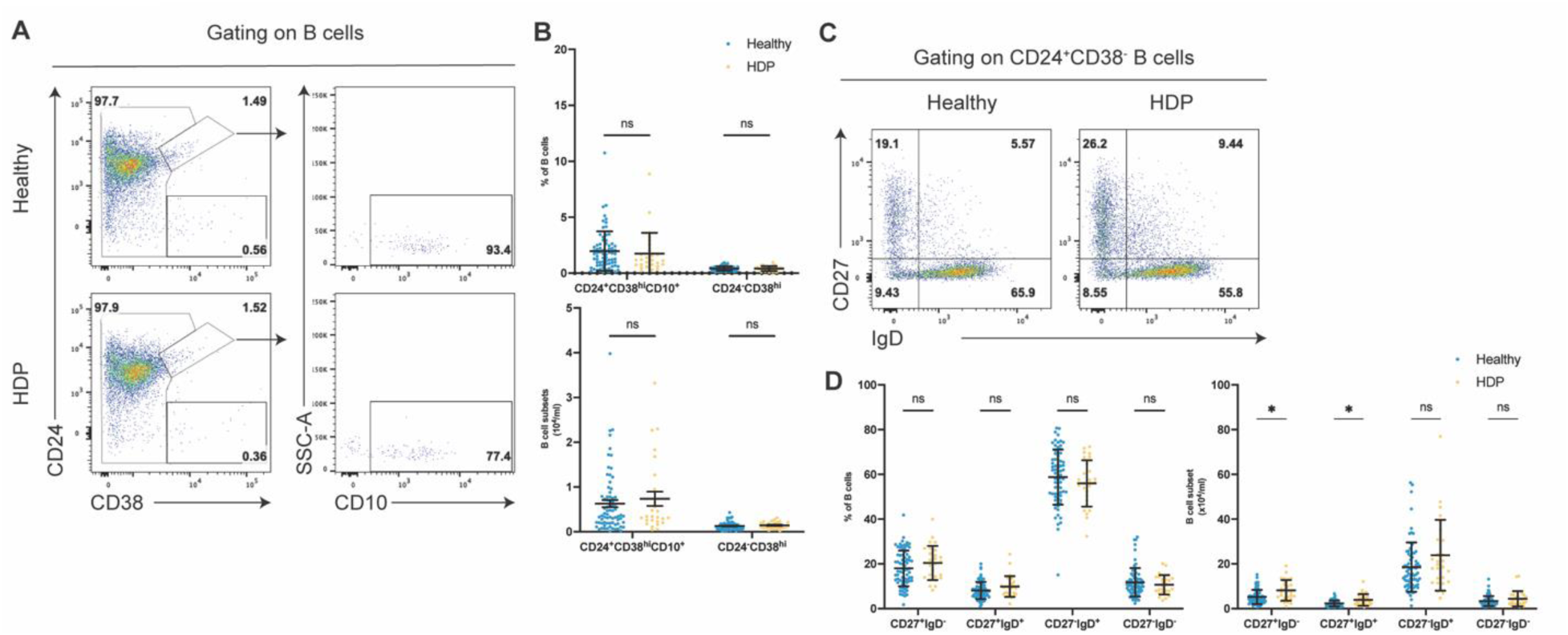
Functional phenotype of B cells in healthy and HDP pregnancies. (A) Representative flow cytometry plots showing CD24^+^CD38^hi^CD10^+^ (transitional B cells) and CD24^-^CD38^+^ (plasmablasts) cells among total B cells. (B) Frequencies and absolute counts of transitional cells and plasmablasts in the peripheral blood of healthy pregnant women and those subsequently diagnosed with HDP (n=73 healthy and n=27 HDP). (C) Representative flow cytometry plots showing CD27 and IgD expression on mature B cells, which are used to define CD27^+^IgD^-^ (class-switched memory), CD27^+^IgD^+^ (non-class-switched memory), CD27^+^IgD^-^ (naïve), and CD27^-^IgD^-^ (double negative memory) B cells. (D) Absolute counts of B cell subsets in the peripheral blood of healthy pregnant women and those who later developed HDP (n=73 healthy and n=27 HDP). Data is shown as mean ± SD.

### Enhanced pro-inflammatory cytokine expression by monocytes preceding the clinical diagnosis of HDP

Monocytes undergo transcriptomic and functional changes during normal pregnancy ^37,39–41^. However, it remains unclear whether these alterations in function are affected in the context of HDP. We therefore examined the functional propensity of monocytes in those with or without a future diagnosis of HDP and focused on cytokine expression following short-term stimulation with lipopolysaccharides (LPS). LPS induces the production of several cytokines in human monocytes, including TNF, IL-6, IL-10, CCL2, and CCL3 (Figure 4A). We found that the absolute counts of TNF^+^ and IL-6^+^ monocytes were elevated in the blood of pregnant women with a future diagnosis of HDP (Figure 4B). Thus, monocytes exhibit pro-inflammatory responses in pregnant women who are later diagnosed with HPD.

**Figure 4.**
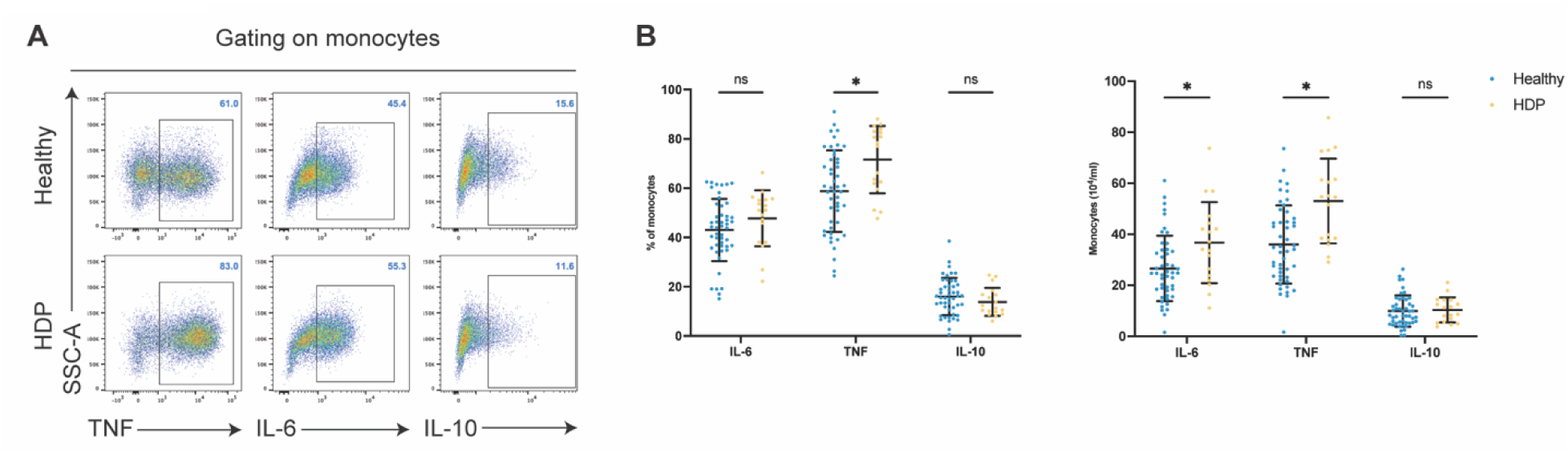
Enhanced pro-inflammatory propensity of monocytes before the clinical diagnosis of HDP. (A) Representative flow cytometry plots showing TNF, IL-6, and IL-10 expression in monocytes following short-term stimulation with LPS. (B) Frequencies and absolute counts of IL-6, TNF, and IL-10 expressing monocytes following stimulation with LPS (n=48 healthy and n=18 HDP). Data is shown as mean ± SD.

## Discussion

In this study, we carried out functional phenotyping of circulating immune cell subsets in the peripheral blood of Black pregnant women obtained during the second trimester to define immune features associated with the future diagnosis of HDP. Our findings identify distinct cellular populations altered months before a clinical diagnosis of HDP. We found elevated counts of T cells in fresh whole blood, and more in-depth functional assays pointed to alterations in particular cellular responses, including enhanced pro-inflammatory cytokine production by CD161^+^ CD4 T cells and monocytes.

Our data indicate that enhanced responses of pro-inflammatory T cells and pro-inflammatory myeloid cells are present before the clinical recognition of HDP. We found that CD161^+^ T cells are more abundant and display an enhanced pro-inflammatory cytokine profile as defined by the expression of GM-CSF and TNF in pregnant women later diagnosed with HDP. GM-CSF typically serves to promote myeloid cell survival and pro-inflammatory functions ^42^. In preeclampsia, increased numbers of pro-inflammatory macrophages are found in the placenta and decidua of patients, and in experimental models, an outcome associated with enhanced GM-CSF expression in decidual tissues ^22,26,27,42,43^. Our finding of an abnormal pro-inflammatory myeloid cell phenotype in the blood of pregnant women later diagnosed with HDP, defined by increased TNF expression, may be related to our observation of increased T cell GM-CSF expression in these patients, since GM-CSF is known to induce expression of several pro-inflammatory transcripts (*TNF*, *IL12B* and *IL23A)* in human monocytes ^44,45^. TNF, in turn, can induce maternal endothelial dysfunction, and its injection or blockade (with etanercept) has been shown to modulate blood pressure and other disease features in pregnant rats ^22^. Our findings extend prior work, including that of Han et al., who identified altered signal pathways across several immune cell populations in pregnant women later diagnosed with preeclampsia. Among these, there was higher basal phosphorylation of STAT5 in the T-bet^+^ subset of effector CD4 T cells ^46^. Phosphorylated STAT5 acts as a positive regulator of effector T cell expression of GM-CSF ^47,48^. Other work identified dysregulated CD4 T cell (Th1 and Th17) responses in the blood following the onset of preeclampsia ^30,31^, and pro-inflammatory CD4 T cells are implicated in the pathogenesis of preeclampsia in animal models ^22^.

We note that anti-inflammatory CD4 T cells and monocytes are not obviously deficient in pregnant women with a future diagnosis of HDP, suggesting that aspects of immunoregulation remain present (but perhaps not enhanced) in the face of the clinical diagnosis. Regulatory CD4 T cells are essential for maternal tolerance to the fetus ^49^, and their frequency increases throughout non-complicated pregnancy ^31^. Alterations in frequency and/or function of regulatory T cells are associated with severe pregnancy-related complications, including preeclampsia, spontaneous preterm birth, and recurrent pregnancy loss ^50^. Our findings of a lack of noticeable change in regulatory CD4 T cell profiles may be consistent with our cohort’s paucity of severe disease. However, we did not sample late pregnancy, and, with our focus on circulating cells, we cannot comment on cellular dynamics within tissues.

Cases had a higher median BMI and increased proportion of obesity compared to controls (Table 1), a well-established clinical risk factor for HDP ^5,51,52^. Our limited sample size precluded rigorous analysis of obesity’s impact on immune signature shifts. However, given the chronic low-grade inflammation associated with obesity ^53^, future studies should investigate whether pregnant individuals with obesity exhibit distinct inflammatory profiles that may contribute to their elevated HDP risk. Longitudinal immune profiling across pregnancy, particularly in obese individuals, may clarify whether specific immune alterations precede or predict HDP development.

Strengths of this study include its prospective cohort design, inclusion of a substantial population of women at highest risk of developing HDP, and the in-depth functional phenotyping of T cells, B cells, and monocytes at a time point preceding the clinical diagnosis of HDP. The prospective design helps to limit inherent biases of case-control studies, and we can infer more about temporality and potential causation than studies assessing immune profiling only at the time of HDP diagnosis. It is understood that racial disparities in HDP are likely driven by social determinants of health, with disproportionate exposure to individual, household, and neighborhood disadvantage among individuals from historically marginalized groups fueling persistent gaps in maternal and infant outcomes ^6^. Future research should explore whether these factors unduly impact immune perturbations, further contributing to the development of HDP. Our work may not be generalizable to a broader population; thus, further investigation is required to ascertain whether our results hold across a racially and ethnically diverse group.

A limitation of this study is the lack of individuals with the more severe phenotypes of HDP, as only 4% of the cohort experienced preeclampsia with severe features (as defined by ACOG ^32^). We cannot establish if a distinct signature is associated with this phenotype of HDP. However, noting that most cases in this study had gestational hypertension, a milder form of HDP, it is noteworthy that such a clear immune signature was appreciated.

In conclusion, our study addresses a current gap in knowledge regarding the involvement of the immune system in the development of HDP in a high-risk and disproportionately affected population. The comprehensive functional immunophenotyping efforts have shed light on functional abnormalities among several cellular populations in this context and have provided the groundwork for further investigation of maladaptive immune responses in more severe cases of HDP (i.e., preeclampsia).

## Perspectives

This study identifies peripheral immune signatures—specifically heightened pro-inflammatory responses among T cells, B cells, and monocytes—in Black pregnant women months before the clinical onset of HDP, offering insights into early disease pathogenesis. These findings suggest that immune dysregulation may precede and contribute to HDP development, rather than arise as a downstream consequence. Immune-based biomarkers assessed in peripheral blood could support risk stratification and earlier identification of at-risk individuals, particularly in high-burden populations. Such profiles may complement emerging molecular tools, including cell-free RNA signatures associated with term and postpartum preeclampsia and gestational hypertension^51^, which are also detectable well before clinical presentation. Earlier detection could enable proactive, individualized interventions to mitigate HDP progression and complications. Current therapeutic strategies—such as low-dose aspirin, antihypertensive therapy, and lifestyle modifications—aim to prolong pregnancy but do not prevent disease onset^32^. Longitudinal studies across gestation, particularly in individuals with obesity or severe HDP phenotypes, are needed to clarify mechanistic pathways and inform the development of targeted therapies. Ultimately, this research reveals immune underpinnings that precede clinical diagnosis of HDP, laying a foundation for biologically informed strategies to reduce maternal morbidity and longstanding disparities in pregnancy outcomes

## Novelty and relevance

a. What is New? Elevation in peripheral T count and increased production of proinflammatory cytokines by CD161^+^ CD4 T cells and monocytes are present months before clinical diagnosis of HDP.
b. What is Relevant? Peripheral pro-inflammatory immune responses in the 2^nd^ trimester of pregnancy provide mechanistic insights into how immune activation may contribute to the pathogenesis of hypertensive processes during pregnancy, which may enable the discovery of biomarkers utilized for earlier detection and intervention before clinical HDP onset. By focusing on Black pregnant individuals, a population disproportionately affected by HDP, we contribute to understanding population-specific risk and the potential role of inflammation in driving disparities.
c. Clinical/Pathophysiological implications? This study identifies a distinct pro-inflammatory immune cell signature in early pregnancy among individuals who subsequently develop HDP, offering potential for early risk stratification.

## Availability of data and material

The anonymized data of this study are available on reasonable request from the corresponding author.

## Acknowledgments

We thank all the patients who participated in this study.

